# Morphology and development of the Portuguese man of war, *Physalia physalis*

**DOI:** 10.1101/645465

**Authors:** Catriona Munro, Zer Vue, Richard R. Behringer, Casey W. Dunn

## Abstract

The Portuguese man of war, *Physalia physalis*, is a siphonophore that uses a gas-filled float as a sail to catch the wind. It is one of the most conspicuous, but poorly understood members of the pleuston, a community of organisms that occupy a habitat at the sea-air interface. The development, morphology, and colony organization of *P. physalis* is very different from all other siphonophores. Here, we propose a framework for homologizing the axes with other siphonophores, and also suggest that the tentacle bearing zooids should be called tentacular palpons. We also look at live and fixed larval and non-reproductively mature juvenile specimens, and use optical projection tomography to build on existing knowledge about the morphology and development of this species. Previous descriptions of *P. physalis* larvae, especially descriptions of budding order, were often framed with the mature colony in mind. However, we use the simpler organization of larvae and the juvenile specimens to inform our understanding of the morphology, budding order, and colony organization in the mature specimen. Finally, we review what is known about the ecology and lifecyle of *P. physalis*.

## Introduction

The pleuston is the floating community of ocean organisms that live at the interface between water and air. This community is exposed to a unique set of environmental conditions including prolonged exposure to intense ultraviolet light, desiccation risk, and rough sea and wave conditions^1^. Despite their tolerance for extreme environmental conditions and the very large size of this habitat, which makes up 71% of the Earth’s surface and is nearly three times the area of all terrestrial habitats, very little is known about the organisms that make up this highly specialized polyphyletic community. One of the most conspicuous, yet poorly understood, members of the pleuston is the siphonophore *Physalia physalis*, commonly known as the Portuguese man of war. The Portuguese man of war is aptly named after a warship: it uses part of an enlarged float filled with carbon monoxide and air as a sail to travel by wind for thousands of miles, dragging behind long tentacles that deliver a deadly venomous sting to fish^2,3^. This sailing ability, combined with a painful venomous sting and a life cycle with seasonal blooms, results in periodic mass beach strandings and occasional human envenomations, making *P. physalis* the most infamous siphonophore^4^.

Siphonophores are a relatively understudied group of colonial hydrozoans. Colonies are composed of functionally specialized bodies (termed zooids) that are homologous to free living individuals. Most species are planktonic and are found at most depths from the deep sea to the surface of the ocean^5–7^. They are fragile and difficult to collect intact, and must be collected by submersible, remotely operated vehicle, by hand while blue-water diving, or in regions with localized upwellings^8,9^. However, *Physalia physalis* is the most accessible, conspicuous, and robust siphonophore, and as such, much has been written about this species, including the chemical composition of its float, venom (especially envenomations), occurrence, and distribution^4,10–20^. Fewer studies, however, have taken a detailed look at *P. physalis* structure, including development, histology of major zooids, and broader descriptions of colony arrangement^21–26^. These studies provide an important foundation for understanding the morphology, cellular anatomy, and development of this pleustonic species. It can be difficult to understand the morphology, growth, and development of *P. physalis* within the context of siphonophore diversity, as the colony consists of highly 3-dimensional branching structures and develops very different from all other siphonophores.

Here, we combine what is already known about morphology and development with new microscopical techniques, including the use of optical projection tomography, and recent phylogenetic and histological knowledge from related siphonophore species, to add new perspectives on the morphology and development of *P. physalis*. As the colony organization is so distinct from other siphonophores, an important first step is to homologize the anatomical axes in developing and mature specimens with other siphonophores. It is then possible to describe the order, pattern, and directionality of budding, and place this within a broader phylogenetic context. There are also still open questions about the homology and origin of some of the unique zooids in *P. physalis*, including the gastrozooid and the tentacle bearing zooid. Additionally, understanding the complex 3D structure of *P. physalis* from written text and hand drawn diagrams can be challenging for a reader that has not spent many hours looking at specimens under a microscope – 3D images and videos can help clarify the complex morphology and arrangement. Finally, we also review what is known about the ecology and lifecycle of this pleustonic species.

## Results and Discussion

### Axes, cormidia, and zooid types

Siphonophores consist of a number of functionally specialized zooids that are homologous to free living polyps or medusae^27^ (Fig. 1). *Physalia physalis* belongs to Cystonectae, a clade that is sister to all other siphonophores^8^. In long-stemmed cystonects (all cystonects except for *Physalia physalis*) gastrozooids (feeding polyps) arise as buds in the anterior of the colony and are carried to the posterior by an elongating stem, while gonodendra (reproductive structures) appear independently along the stem^28^. In all cystonects, the gonodendra are compound structures, containing gonophores (reduced medusae, bearing a gonad), palpons (derived gastrozooids, that lack tentacles in cystonects), and nectophores^27^. *P. physalis* gonodendra have these zooids, as well as ‘jelly polyps’ that are reduced nectophores of unclear function^22,26^. Cystonects are dioecious, and all the gonodendra in a colony bear gonophores of only one sex.

**Figure 1:**
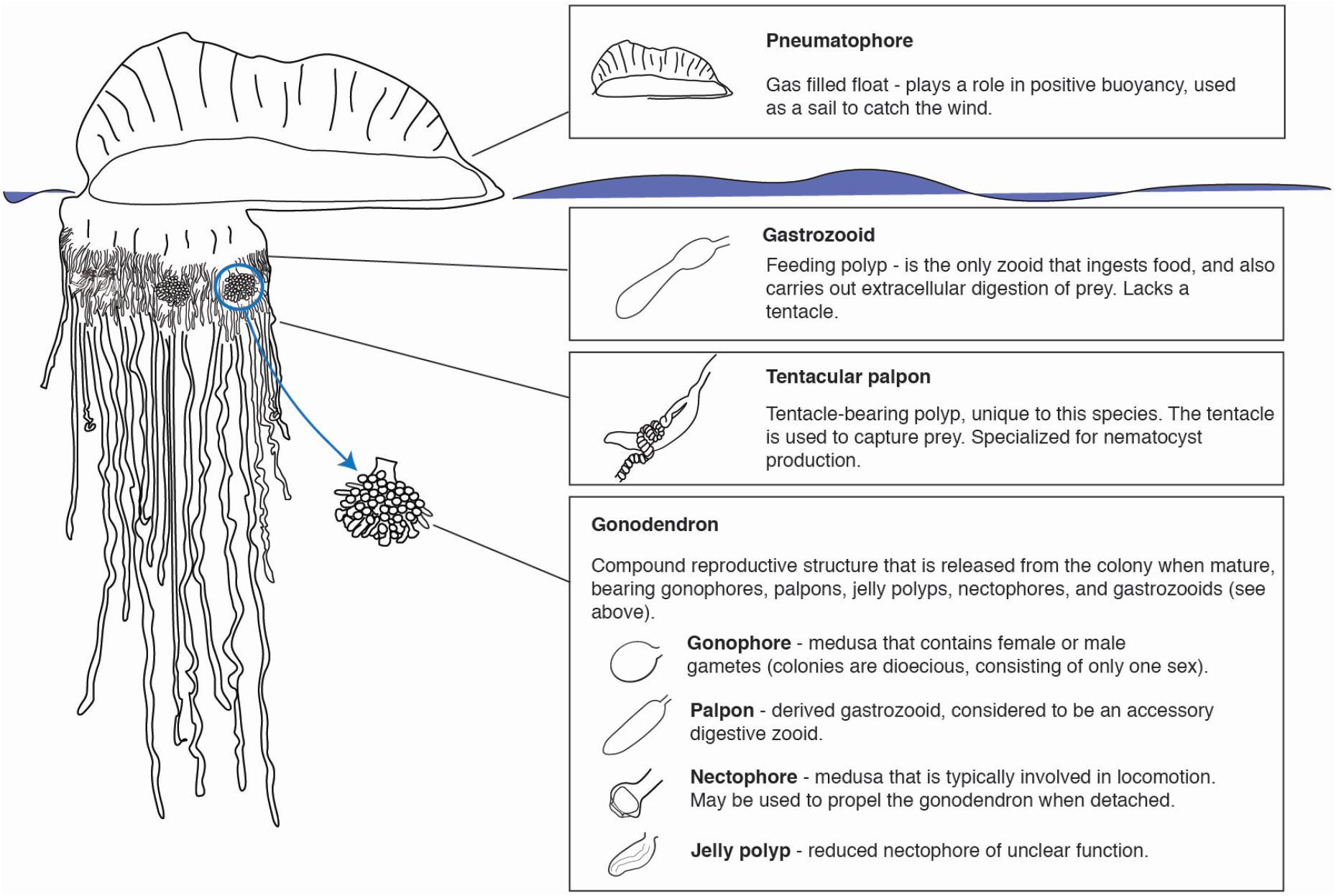
Schematic showing the anatomy of a *Physalia physalis* colony, with descriptions of the function of each zooid.

As compared to other siphonophore species, including other cystonects, *Physalia physalis* is peculiar with regards to its colony organization (Fig. 1). *P. physalis* is also the only siphonophore species where the gastrozooid, the primary feeding zooid, does not have a tentacle for prey capture. The only exception is the protozooid (the first gastrozooid to form during development), which is essentially a typical siphonophore gastrozooid, with a mouth, tentacle and small basigaster region^26,29^. Apart from the protozooid, in *P. physalis* the tentacle is borne on a separate zooid that Totton called the ampulla^26^. Other authors refer to either the zooid or the attached tentacle as a dactylozooid^10,13,21,30^ - the term dactylozooid has historically been applied to palpons in other siphonophore species but is not currently used, and dactylozooids are specialized palpon-like defensive zooids in other hydrozoans^31–33^. To avoid confusion about the homology of this zooid, we suggest that the term dactylozooid should not be used, as we consider this zooid to have arisen *de novo* in *P. physalis* and is not likely homologous to dactylozooids in other hydrozoans. Additionally, the term ampulla is also commonly associated with the terminal vesicle of the tricornuate tentillum of agalmatids^27^. We favor reviving Haeckel’s ‘tentacular palpon’ to refer to this zooid^34^, which not only has precedence, but also matches the likely hypothesized origin of this zooid (see below).

Haeckel outlined two possible hypotheses for the origin of tentacular palpons - the first hypothesis, promoted by Huxley, is that they are not zooids, but are instead secondary diverticula at the base of the tentacle that function similarly to ampullae in echinoderm tube feet^34,35^. In the second hypothesis, modification and subfunctionalization of an ancestral gastrozooid gave rise to two separate zooids - a gastrozooid without a tentacle and a tentacular palpon with a tentacle. Totton proposed a modification of the first hypothesis, and suggested that the ‘ampulla’ is a hypertrophied basigaster (aboral region of a gastrozooid that plays an active role in nematogenesis) that has separated from the remainder of the gastrozooid^26^. However, we favor the second hypothesis, based on observations of the gastrozooid and tentacular palpon (Figs. 3,4, 7, 6; supplementary videos 1, 2). The gastrozooid and tentacular palpon are borne on separate peduncles (Figs. 7A, 6A, 6B; supplementary video 2), and develop from distinct, separate buds (Figs. 3A, 3B, 4; supplementary video 1). Thus, the tentacular palpon is a derived gastrozooid, unique to *Physalia physalis*, that has an enlarged tentacle, no mouth, and is functionally specialized for nematocyst production. The gastrozooids in *P. physalis* are also derived gastrozooids that have lost tentacles and are functionally specialized for feeding only. The subfunctionalized gastrozooid hypothesis is also more parsimonious than the other hypotheses, as the modification and subfunctionalization of zooids is common in siphonophores - palpons, for example, are considered to be derived, modified gastrozooids that typically have a reduced tentacle^27^.

Historically, there was no consistent terminology to describe the axes of mature siphonophore colonies. A standardized scheme was developed to describe mature planktonic siphonophore colonies, with the anterior end of the colony as the pneumatophore and the posterior end of the colony as the oldest^36^ (Fig. 2A). The dorsal-ventral axis is perpendicular to this axis, with siphosomal zooids attached to the ventral side of the stem. Left and right are determined as perpendicular to the anterior-posterior and dorsal-ventral plane. The oral end of the larva corresponds to the posterior of the mature colony^36^. As *Physalia physalis* is a pleustonic species, with distinctive colony morphology and arrangement, it is important to homologize the axes with other siphonophores. Totton does not use the terms anterior-posterior, and defines an oral-aboral axis that corresponds directly to the larval axis, with the protozooid, the first feeding zooid (Fig. 2B), on the oral end and the apical pore (the pore is the site of invagination forming the pneumatophore) of the pneumatophore on the aboral end^26^. The oral end of the colony thus corresponds to the posterior end^36^. This corresponds directly with the anterior-posterior axis defined by other authors^25,35^, with the apical pore defined as the anterior of the colony. To keep terminology consistent across all siphonophores, we will follow this convention, with the anterior corresponding to the apical pore and the posterior corresponding to the protozooid (Fig. 2B). The dorsal-ventral axis is perpendicular to this plane, with the dorsal side towards the crest of the float and zooid attachment on the ventral side (Fig. 2C). We will follow the same left-right and proximal-distal axis conventions. While zooid attachment is on the ventral side, there are very clear left-right asymmetries in the placement and growth of zooids in this species, and colonies are either left-handed or right-handed.

**Figure 2:**
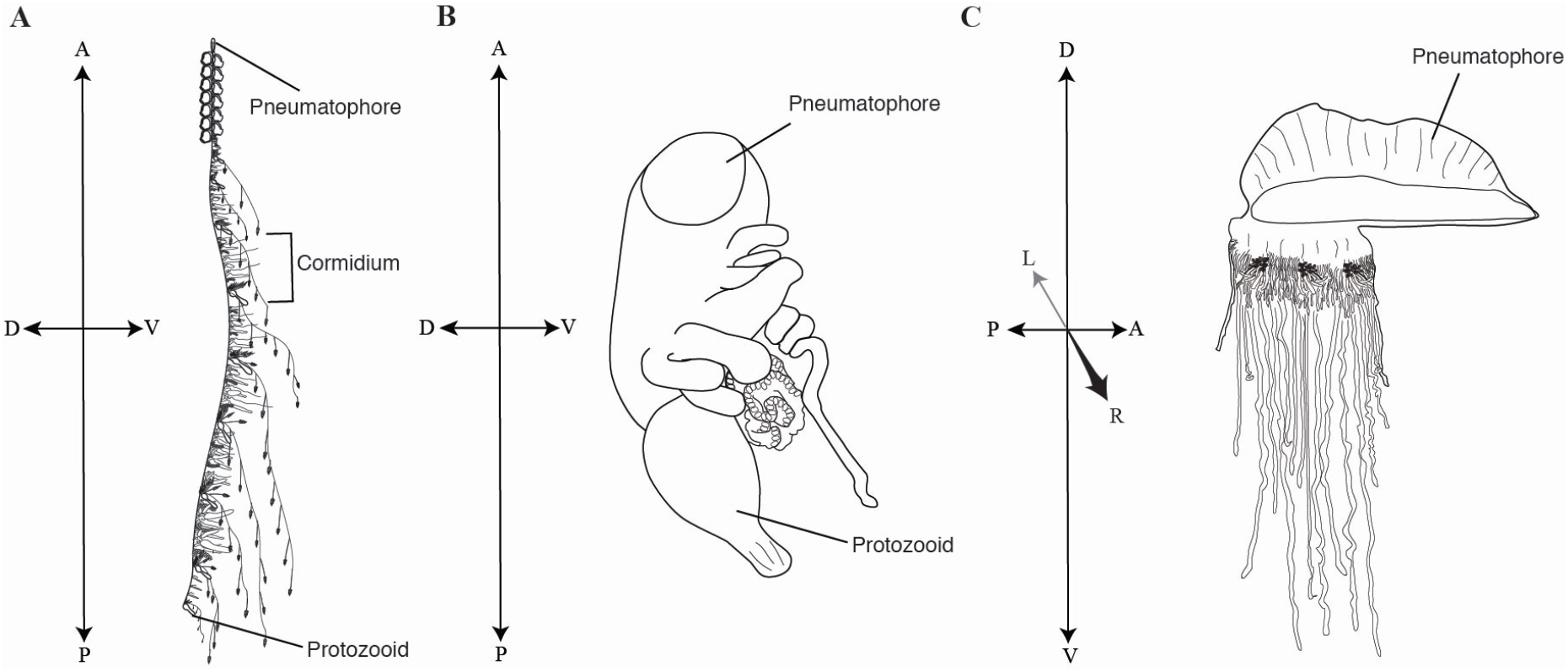
Colony orientation in siphonophores. A - anterior, P - posterior, D - dorsal, V - ventral, L - left, R - right. A. Schematic of a mature colony of the siphonophore *Nanomia bijuga*. Drawing by Freya Goetz, wikimedia commons. B. Schematic of a developing *Physalia physalis* larva. Drawing based on photograph by Linda Ianniello. C. Schematic of a mature *Physalia physalis* colony.

Cormidia are typically defined as a group of zooids that are reiterated along the siphosomal stem in many siphonophore species^27^ (Fig. 2A). Many authors refer to ‘cormidia’ in *Physalia physalis*. Cystonectae, the clade to which *P. physalis* belongs, are sister to Codonophora^6,8^. Cystonects produce all zooids from single buds that arise along the stem, while probud subdivision (all zooids in a cormidium arise from a single bud) is a synapomorphy of Codonophora^28^. Probud subdivision is associated with the origin of cormidia along the branch that leads to Codonophora^28^. Due to this, and the fact that *P. physalis* has very distinct development and morphology, we will not apply the term ‘cormidia’ to describe *P. physalis* organization.

### Larval development and morphology

Larval development has not been observed directly, and development has been described by comparing the morphology of fixed specimens^23,24,26^. The smallest described larva is 2mm, and consisted of a pneumatophore and a developing protozooid with a tentacle^26^. The pneumatophore forms in a manner similar to other siphonophores, with an invagination of the aboral end of planula forming the pneumatosaccus^24,29,37^ (Fig. 5). Okada suggests that the apical pore that is formed by this invagination is completely closed in larval *Physalia* (float length 2mm) and controlled gas release from the pneumatophore, as in some other siphonophore species, is no longer possible^23^. However, Mackie suggests that the pore is not completely closed even in mature colonies, but the pore is so tightly constricted that gas release is unlikely to occur naturally^22^. Other reports suggest that young *Physalia* may be able to release gas from the pore^38^. In the earliest stages, there is no separation between the gastric cavity of the protozooid and the main gastric cavity^24^. The pneumatosaccus, that is formed via the invagination, protrudes into the main gastric cavity and is connected at the site of invagination^24^. As the protozooid differentiates, a septum separates the gastric cavity of the protozooid from the main gastric cavity^23^.

Anterior to the protozooid, three buds arise on the ventral side as three transverse folds^26^. Based on our observations of the budding order and the relative size of the zooids, the posterior most of these three buds is a gastrozooid G1, followed by a second gastrozooid G2 and tentacular palpon (labelled Tp1) (Figs. 3A, 3B, 4). The observation of the three transverse folds suggests that G1,G2, and Tp1 all arise at the same time^26^. The third gastrozooid (G3) subsequently appears anterior to gastrozooids G1, G2, and tentacular palpon Tp1. Totton numbers the buds based on the hypothesized ‘cormidia’ to which they belong in the mature colony, but not based on their order of appearance^26^. Okada numbers the buds based on hypothesized order of appearance, which differs from ours only in that G2 is considered the first bud, perhaps based on size, and G1 is considered the second^23,24^ (Figs. 3A, 3B). The gastrozooid labelled G2 here is larger in older specimens (Figs. 3C, 3D, 4), but not in the youngest developing specimen (Fig. 3A).

**Figure 3:**
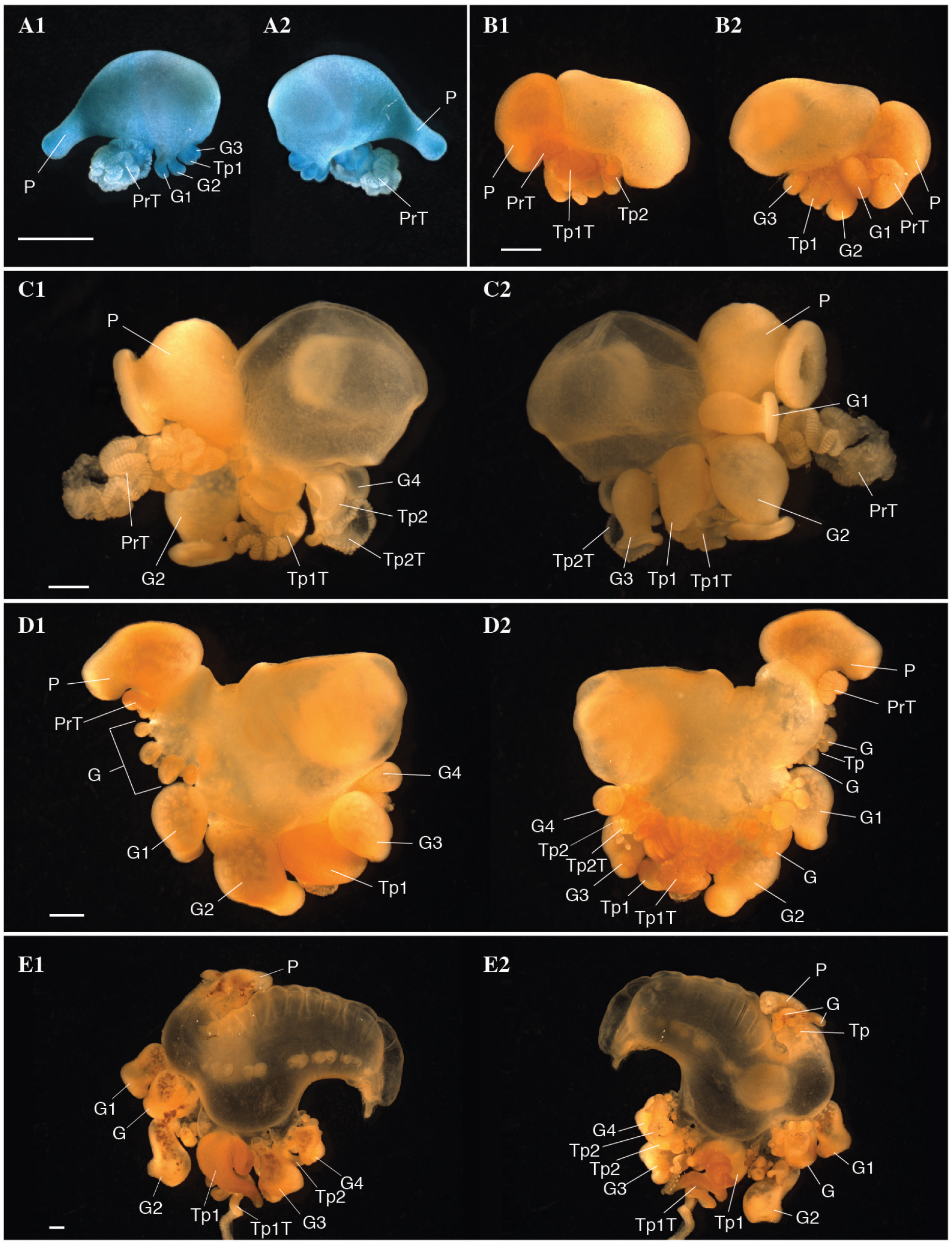
Photographs of formalin fixed developing *Physalia physalis*. Five different specimens are shown in A-E. Photographs 1 and 2 represent left/right sides of the same specimen. Scale bar is 1mm. Abbreviations: Tp: Tentacular palpon (number indicates order of appearance); G: Gastrozooid (number indicates hypothesized order of appearance; G1 and G2 likely appear at the same time) P: Protozooid; PrT - Tentacle of protozooid; TpT: tentacle of tentacular palpon (number indicates order of appearance).

*Physalia physalis* colonies can be either left or right handed, and the location of first tentacular palpon (Tp1) and the attachment point of the tentacle is the first indicator of left-right asymmetry^24,26^. The tentacle of the tentacular palpon is placed either on the left or right side, depending on the handedness of the colony (Fig. 3, 4; supplementary video 1). The secondary series of buds always appear on the same side as the tentacular palpon tentacle. The attachment point of the tentacle of the protozooid may even be an earlier indication of left-right asymmetry (Fig. 3, 4; supplementary video 1). As live embryos are not available, it remains an open question as to whether left-right asymmetries are established via molecular mechanisms similar to those underlying left-right asymmetry in bilaterians^39^.

**Figure 4:**
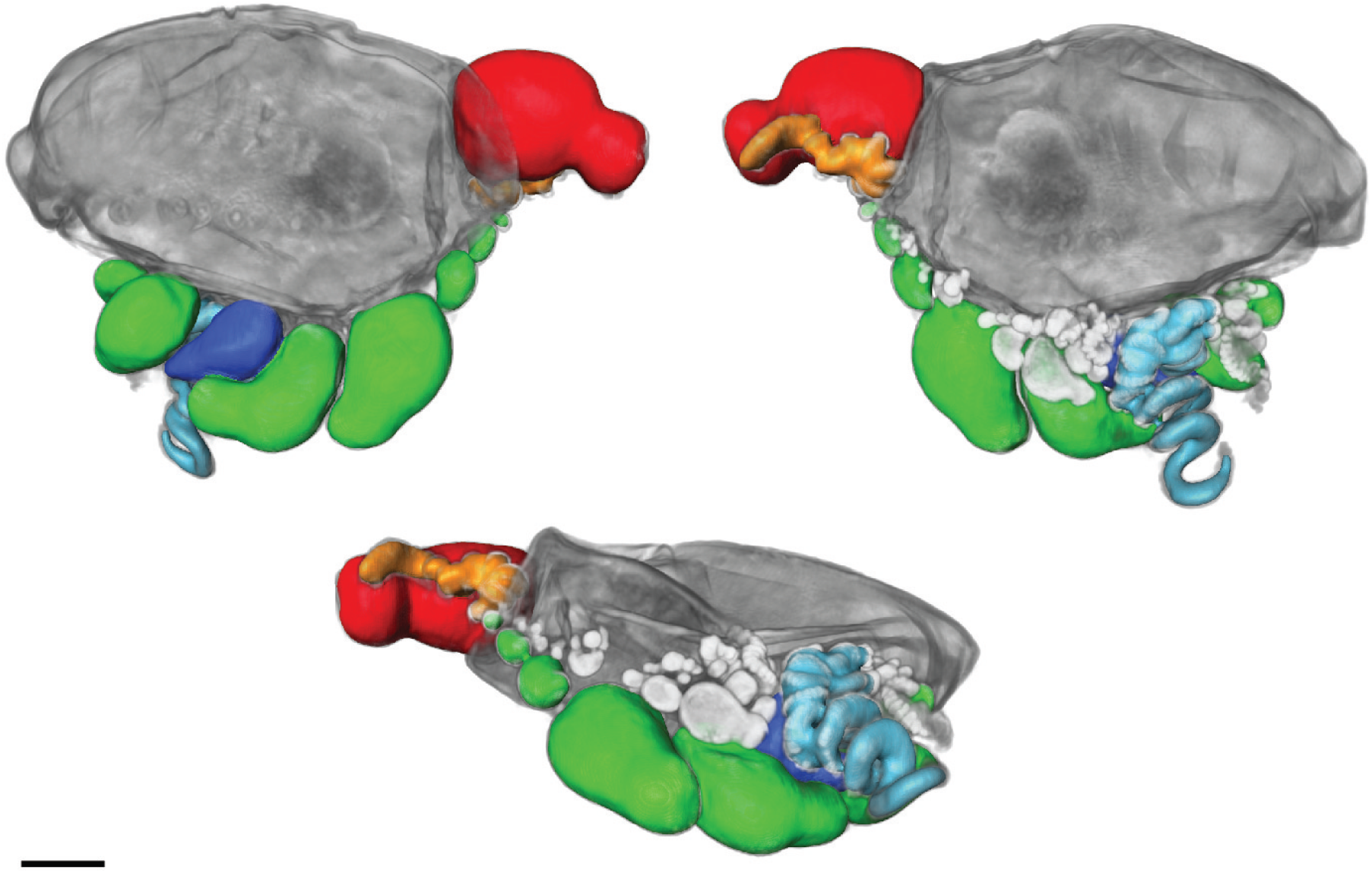
Images of formalin fixed larval *Physalia physalis*, images obtained by optical projection tomography. Images are different views of the same specimen. Scale bar is 1 mm. The 3D image was segmented and false-colored to highlight different morphological features. Green- gastrozooids; Red- Protozooid; Orange- tentacle associated with protozooid; Dark blue- Tentacular palpon; Light blue- tentacle associated with tentacular palpon. Gastrozooids and tentacular palpons forming at the base of the first set of gastrozooids and tentacular palpon are unlabelled and are light grey in color.

As the organism grows and the pneumatosaccus expands anteriorly, new tentacular palpons grow at the base of the original gastrozooids (Figs. 3B2 and 4). In larger specimens, new gastrozooid and tentacular palpon buds form anterior and posterior to the three gastrozooids (G1, G2, G3) and tentacular palpon (Tp1) (Figs. 3C1, 3D). A secondary series of buds also form at the base of the gastrozooids in line with the first tentacular palpon (either left or right, depending on the handedness of the colony)^23^ (Figs. 3D2, 4). Additionally, in the expanding space between the protozooid and the primary series of gastrozooids, a series of buds form (Fig. 3D, labelled “G”; Fig. 4, gastrozooids (in green) closest to protozooid). This region of growth directly anterior to the protozooid (Figs. 3D, labelled “G”) is distinguished from the original region by Totton as the “oral zone”, while the original series of buds (including G1, G2, G3, G4, Tp1, Tp2, and secondary buds) are the “main zone”^26^. To keep naming consistent with the axes, we propose calling the oral zone the “posterior zone”. In older larvae the protozooid and posterior growth zone are physically separated from the main zone, due to elongation of the stem/float carrying the posterior growth zone away from the main growth zone (Fig. 3E).

As *Physalia physalis* continues to grow, new space along the ventral side in the main zone is occupied by new buds in line with the original series of gastrozooids (G1, G2, G3 etc.) and tentacular palpon. Additional secondary clusters of buds also continue to arise both in the posterior and main zone, either to the left or right according to the handedness of the colony (Fig. 4). A crest begins to become visible (Figs. 3D, 3E), and the float expands. Once the float is fully expanded, and the colony is floating on the ocean surface, branching and growth begins to occur in the dorsal-ventral plane (Fig. 2C). In fully mature specimens, zooids occupy the space between the posterior and main zones, and the gap (termed the basal internode) between the two zones of growth is not visible.

Superficially, the series of buds in the posterior zone resembles the growth zone of related species, such as *Nanomia bijuga*^40,41^. We do not know the order of bud appearance, however based on the relative size of the gastrozooids (Fig. 4), new buds in the posterior zone appear to arise posterior-anterior along the ventral side in an inverse direction to other siphonophore species (Fig. 2B). This does not fit with the definition of axes as defined by Haddock *et al*, with buds arising in the anterior and being carried by elongation of the stem to the posterior^36^. Patterns of growth are very different in *Physalia physalis*, however this may suggest that during early development growth patterns are inverted in this species. According to our numbering system, the original series of buds (G1, G2, Tp1) also arise posterior-anterior, although subsequent buds in the main zone arise both anterior and posterior to these zooids.

The patterns of growth that can be observed from fixed developing *Physalia physalis* specimens suggests that while there are many similarities between this species and other siphonophores, there are many differences that are unique to this species. In other siphonophore species, ontogenetic series of zooids are arranged linearly along a stem with the oldest at the posterior and the youngest in the anterior^28,41^, although new zooids are observed to arise along the stem in some species^42^. In *P. physalis* there are three major axes of growth – along the ventral side, posterior-anterior in the posterior growth zone (Fig. S5A), as well as anterior and posterior of the main zone; secondary buds to left or right of the original series of buds along the ventral side, depending on the handedness of the colony; and finally in mature specimens, growth proceeds proximal-distal from the ventral side (Fig. S5B).

### Morphology and zooid arrangement of mature *Physalia physalis*

Juvenile (sexually immature) and mature *Physalia physalis* float on the ocean surface with the pneumatophore, or float, above and on the surface of the water and all of the zooids are below the water surface. In juvenile *P. physalis* the pneumatophore will continue to grow in size, but it resembles the fully mature form. As in other siphonophores, the pneumatophore is a multi-layered structure, consisting of an outer codon, a pneumatosaccus, and a gas gland^22^ (Fig. 5). The outer codon consists of ectoderm, mesoglea, and endoderm^22^. Within the codon is the pneumatosaccus, formed by invagination, consisting of endoderm, mesoglea, ectoderm, a chitinous layer secreted by the ectoderm, and the gas space^22^. At one end of the pneumatosaccus is an expanded layer of ectodermal cells that form the gas gland^11,22^. Aeriform cells within the gas gland produce carbon monoxide to fill the float, however the percentage of carbon monoxide within the float is lower than other siphonophores due to diffusion and gas exchange^14,17,43^. Unlike other siphonophores, the pneumatophore is greatly expanded, and the pneumatosaccus is free within the gastric cavity and attached only to the site of invagination at the anterior of the colony^22^. Dorsal processes of the pneumatosaccus fit into pockets of the crest of the codon, and muscular contractions of the codon enable the pneumatosaccus to expand into this space and erect the sail – this ‘pneumatic skeleton’ is likened to a hydrostatic skeleton^22^. The zooids are all attached on the ventral side (displaced either to the left or right) and share this common gastric cavity – this region is likely homologous to the stem of other siphonophores^44^.

**Figure 5:**
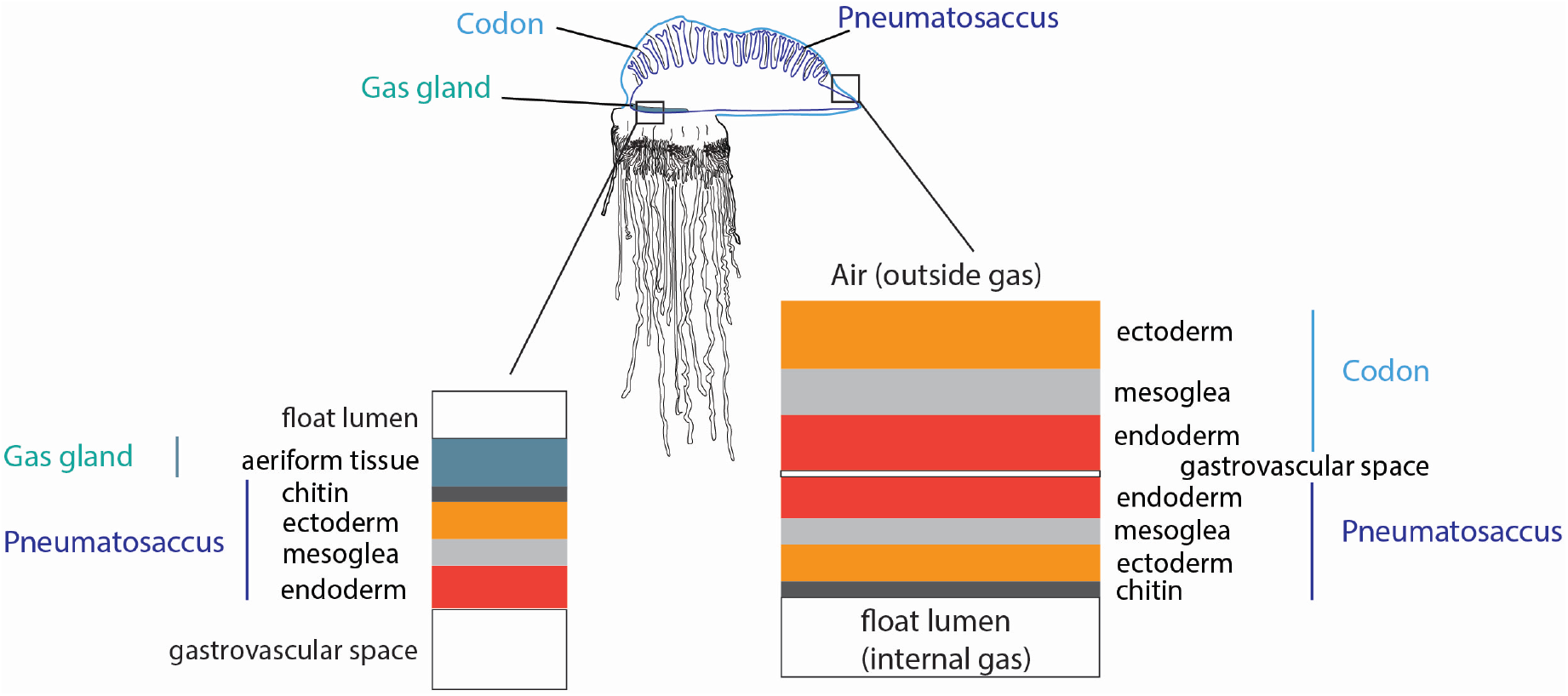
Schematic of the cell layers in the pneumatophore, showing the distinction between the codon, pneumatosaccus, and gas gland. Orange - ectoderm; dark grey - chitin; red - endoderm; light grey - mesoglea.

In juvenile *Physalia physalis*, projections extend from the ventral ‘stem’, carrying zooids distally away from the float. The colony arrangement of *P. physalis* appears crowded and lacking in structure, particularly in fully mature specimens, however there is a distinct pattern of growth. The best descriptions of colony arrangement in mature specimens are given by Totton, who suggested that growth occurs through the formation of tripartite groups^26^ (Figs. 7, 6; supplementary video 2). The tripartite groups consist of a tentacular palpon with an associated tentacle, a gastrozooid, and a gonodendron at the base of the gastrozooid^26^. The morphology of *P. physalis* is clearest in juvenile specimens, where the gonodendron is not fully developed and developing tripartite groups are easily identifiable (Figs. 6, 7B). The gonodendron is a structure that consists of a number of different zooids, including gastrozooids, male or female gonophores (colonies are dioecious, and as such, colonies are either male or female), nectophores, jelly polyps, and also palpons.

**Figure 6:**
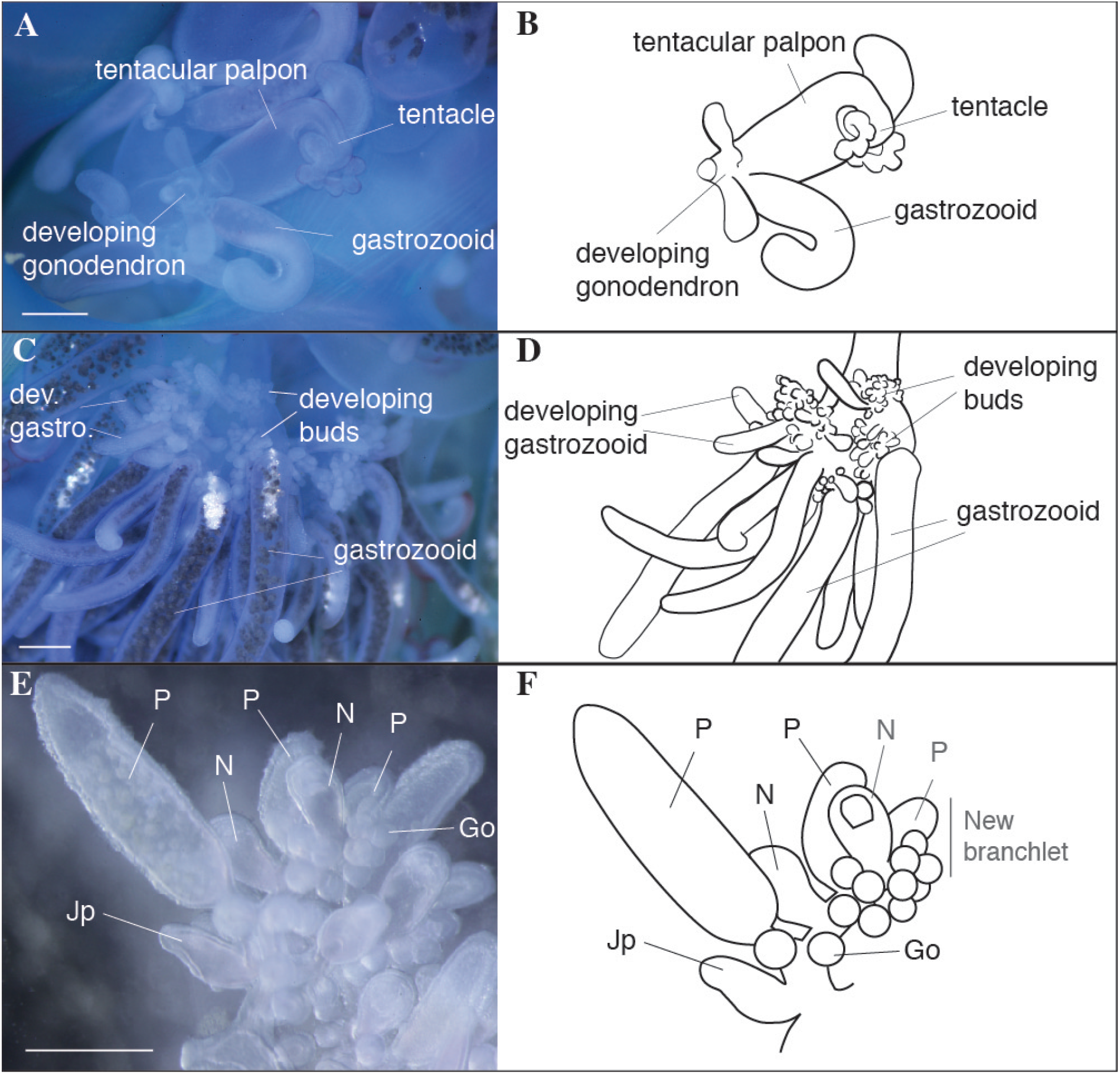
Photographs of live juvenile *Physalia physalis*. Scale bar is 1mm. A. Developing tripartite group, with gastrozooid, tentacular palpon and developing gonodendron. B. Schematic of the tripartite group. C. Developing gonodendron with mature gastrozooids and buds that will give rise to gonophores, nectophores, palpons. D. Schematic of the developing gonodendron. E. Close up of branchlet within the gonodendron, from proximal to distal: jelly polyp (Jp), palpon (P), nectophore (N), palpon, with gonophores (Go) along the branchlet, additionally there is a nectophore, palpon and gonophores that are part of a new branchlet. Schematic of a close up of a branchlet within the gonodendron.

**Figure 7:**
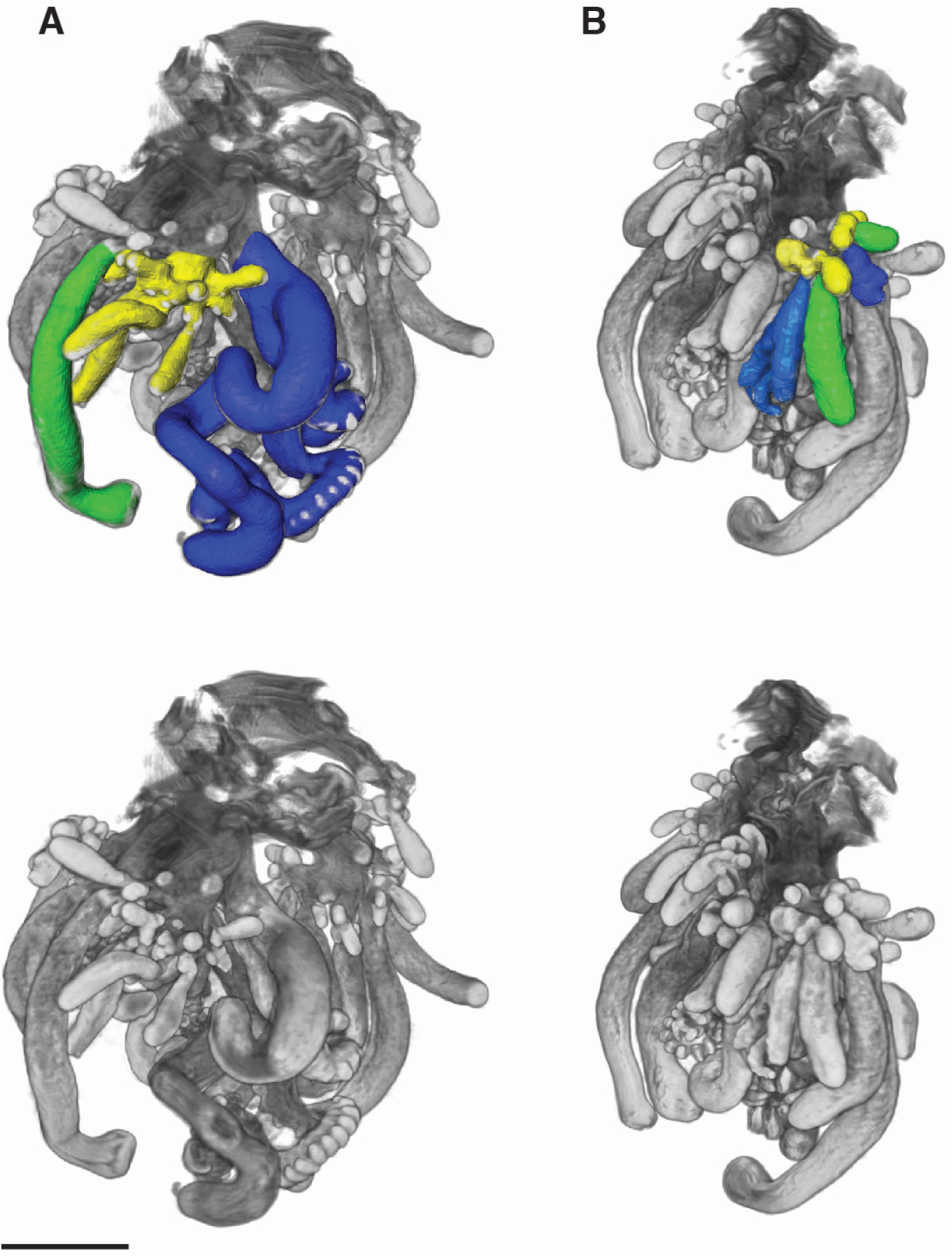
Images of formalin fixed juvenile *Physalia physalis* zooids, images obtained by optical projection tomography. Images are different views of the same specimen. Scale bar is 2mm. The 3D image was segmented and false-colored to highlight tripartite groups. The un-segmented image is shown below. Green- gastrozooid; Dark blue- tentacular palpon; Yellow- developing gonodendron. A. Tripartite group with developing tentacular palpon, gonodendron and gastrozooid. B. Two sets of developing tripartite groups at different developmental stages are highlighted, while others are visible but not segmented.

Tripartite groups are carried down by elongated projections of the stem, with successive tripartite groups forming at the base of the older groups. In mature colonies, the oldest zooids are located distally, with developing zooids in tripartite groups forming proximally to the float (Fig. S5B). The exception to this appears to be the very oldest zooids that form during early development (Fig. 3, 4), that remain attached proximally to the stem via long peduncles. There are differences in the rate of growth and appearance of zooids in the tripartite groups: the tentacular palpon and gastrozooids both develop precociously, while the gonodendron develops and matures later^26^. The developing and mature tentacles could be distinguished not only by size and length, but also by color – the tentacles of the mature tentacular palpons are a turquoise blue, while the buttons of the developing tentacle are a purple/pink color. The blue pigment of *P. physalis* is suggested to be a bilin-protein complex, and the green, purple, and pink coloration in other tissues are caused by unconjugated bile pigments, which are likely sourced from their diet^12^.

The gonodendra are highly complex branching structures. We are not able to build much upon the description by Totton^26^ of the structure of the gonodendron, but we do attempt to simplify aspects of his description here, based on our observations. In the juvenile specimens, we were able to observe developing gonodendra with mature gastrozooids (what Totton calls ‘gonozooids’, or secondary gastrozooids) and clusters of buds at their base that will subdivide and give rise to all the other zooids within the gonodendron^26^ (Fig. 6B). The peduncles at the base of the gastrozooids form the major branches within the gonodendron^26^. Branching can be observed at two levels: major branches formed by the peduncle of the gastrozooid (Fig. 6B); and branching structures at the base of the gastrozooids, that are formed by probuds (Fig. 6B “developing buds”) that subdivide, branch and re-branch, and form a series of branchlets along which nectophores, jelly polyps, palpons, and gonophores form (Figs. 6E,6F). The branchlets of the gonodendra typically consist of a series (proximal to distal) of a jelly polyp and more developed palpon, followed by a nectophore and palpon, with ~ 10 or more male or female gonophores (depending on the sex of the colony) forming along the branchlet (Figs. 6E,6F). Totton refers to the section with the jelly polyp and palpon as the terminal section of the branchlet^26^, while the sub-terminal portion of the branchlet may become a palpon and nectophore (Figs. 6E,6F), or continue dividing into a new terminal and subterminal portion. New probuds form in the region directly opposite the location of jelly polyps, giving rise to new branchlets, that in turn re-branch opposite the location of the jelly polyp^26^. Sometimes a branchlet can consist only of a palpon and jelly polyp^26^.

### Ecology and lifecycle

*Physalia physalis* is a cosmopolitan species, found in tropical and subtropical regions of the ocean, as well as occasionally in temperate regions^26^. Historically, a large number of *Physalia* species have been described on the basis of size, color, and location^26,35,44,45^, however, there is currently only one recognized species of *Physalia* – *P. physalis*^27^. The different species that have been identified are suggested to be different developmental stages^23,26,27^. However, nothing is known about genetic diversity among populations of *P. physalis* in the Atlantic or the Pacific/Indian Ocean. One local study has been conducted, using two genetic markers, that showed substantial genetic diversity among *Physalia* off the coast of New Zealand^46^, however global studies using more markers would help clarify whether this reflects intra-specific genetic diversity or if there is cryptic diversity.

As larval development has not been observed directly, everything that is known about the early stages of this species is known from fixed specimens collected in trawl samples^23,24,26^. Gonodendra are thought to be detached by the colony once they are fully mature, and the nectophores may be used to propel the gonodendron through the water column^25,26^. Released mature gonodendra have not been observed, and it is not clear what depth range they occupy^25,26^. It is also not known how the gonodendra from different colonies occupy a similar space for fertilization, or if there is any seasonality or periodicity to sexual reproduction. Embryonic and larval development also occurs at an unknown depth below the ocean surface^26^ (Fig. 8). After the float reaches a sufficient size, the juvenile *P. physalis* is able to float on the ocean surface.

**Figure 8:**
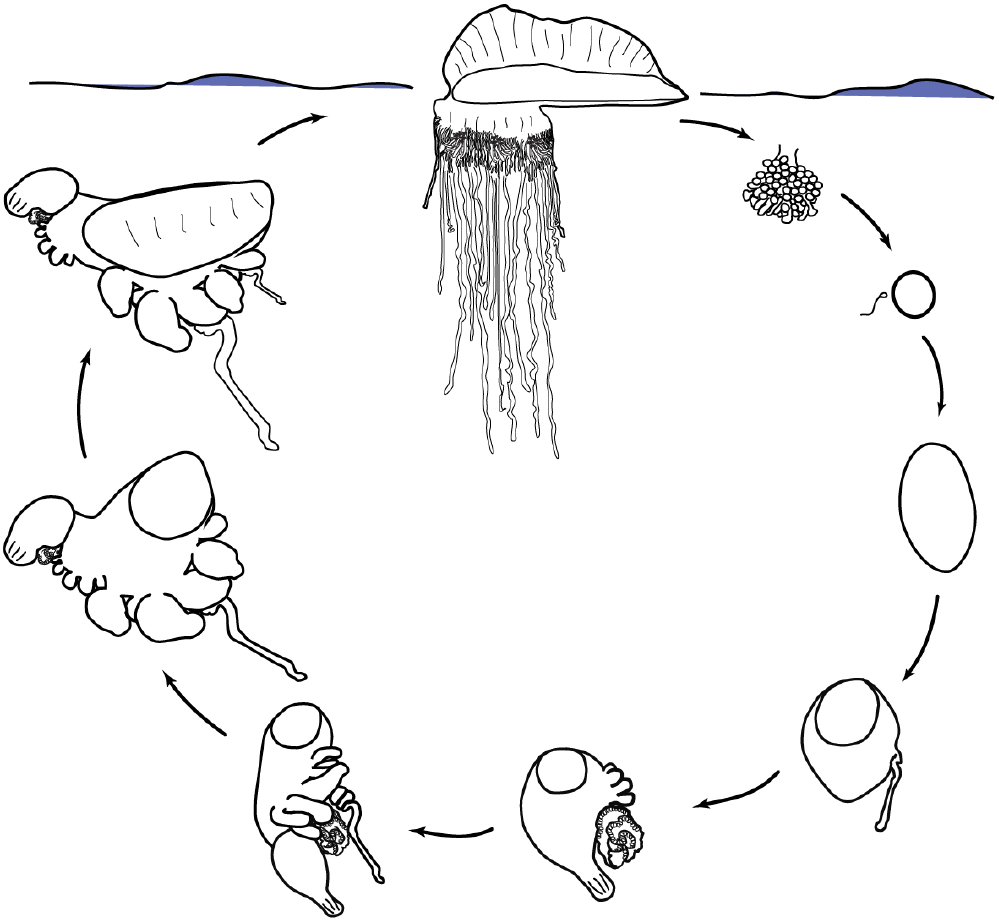
Schematic of the lifecycle of the Portuguese man of war. The mature *Physalia physalis* is pictured floating on the ocean surface, while early development is thought to occur at an unknown depth below the ocean surface. The gonodendra is thought to be released from the colony when mature. The egg and planula larva stage have not been observed. The egg and planula drawings are from a *Nanomia bijuga* lifecycle schematic drawn by Freya Goetz, wikimedia commons.

Mature *Physalia physalis* uses its sail to catch prevailing winds. Muscle contractions of the outer codon of the pneumatophore force increased pressure within the pneumatosaccus and enable the erection of the crest^22^. This is the only known active contribution to locomotion – the nectophores within the gonophore are not thought to play any role in active propulsion of the colony (although they may play a role once the gonodendron is released)^26^. The alignment of the sail relative to the wind (left-right handedness), is established during early development, and while it has been suggested that left-handed colonies are dominant in the Northern Hemisphere as a result of prevailing winds, and right-handed colonies are more prevalent in the Southern Hemisphere^19,47^, there is no evidence to support this^15,26^. Wind fluctuations are likely to result in random distribution of both forms regardless of hemisphere, although strong sustained winds from the same direction do appear to result in the stranding of a particular type^15,48^. Totton suggests that left-right asymmetry is established by the prevailing wind on the first windy day^26^, however this is unlikely, as the asymmetry is present early in developing specimens.

The tentacles of the Portuguese man of war can reach up to 30m in mature colonies, and are dragged through the water due to the wind, adhering to fish and fish larvae that they encounter. Fish and fish larvae comprise 70-90% of their diet, and the nematocyst batteries on the tentacles of *Physalia physalis* contain a single type of nematocyst that is only able to penetrate soft bodied prey^49,50^. The nematocyst delivers a toxin that leads to hyperventilation, immobilization and, in high doses, death^51^. Once a tentacle comes into contact with its prey, the prey is carried up towards the gastrozooids near the base of the float. The gastrozooids respond immediately to the capture of prey, and begin writhing and opening their mouths^52^. Many gastrozooids attach themselves to the prey – upwards of 50 gastrozooids have been observed to completely cover a 10cm fish with their mouths spread out across the surface of the fish^16^. The gastrozooids release proteolytic enzymes to digest the fish extracellularly, and are also responsible for intracellular digestion of particulate matter^22,53^. The digested food products are released into the main gastric cavity for uptake by the rest of the colony^22,53^. While *P. physalis* is a voracious predator of fish, it is predated upon by sea turtles^54,55^, and *Glaucus atlanticus* and *Glaucus marginatus*, species of nudibranch that store intact *Physalia* nematocysts and redeploy them for their own defense^56–58^. A number of juvenile fish live commensally with *Physalia* and are found near the gastrozooids and gonodendra, however one species, *Nomeus gronovii*, has been observed to swim among and feed upon the tentacles^30,59^. *Nomeus gronovii* is significantly more tolerant of *Physalia* venom than other species, but can nevertheless be killed by *P. physalis*^26,60^.

## Conclusions

*Physalia physalis* differs significantly from all other siphonophores in terms of its habitat, development, body plan, and colonial organization. The radical modification of the colony body plan is likely associated with a transition from a planktonic to pleustonic lifestyle. Using photographs, specimens and new volumetric imaging methods to create 3D reconstructions, we were able to clarify aspects of *P. physalis* colony organization in juvenile specimens, and also early development in larval specimens. The study underscores the value of fixed specimen collections – all of the developing specimens used in this study were collected in the 1970s and 1980s, and it was still nevertheless possible to 3D image these colonies using fluorescent stains. Optical projection tomography is particularly useful for imaging these complex, highly branching structures, and we are able to use these images to build upon the existing knowledge about the development, morphology and colony organization of this species. In particular, larval and juvenile specimens were key for this work, because growth and secondary budding in mature specimens makes it significantly more difficult to understand the order and pattern of growth.

Many open questions remain about this species, however. While Totton was able to observe gonodendra that are more mature than those examined in this study, fully mature gonodendra with mature eggs or sperm have not been described yet. Mature gonodendra are hypothesized to be released into the water column, however there is no data on the depth ranges that the gonodendra occupy. Additionally, there is also no information about the depth at which any of the early developmental stages can be found, nor their ecology. While there is abundant data on the occurrence and location of *P. physalis*, particularly beached specimens, there is frequently little recorded information about the size of the colony, and it is not clear if there is seasonality to their reproduction. Most of our experiences of the Portuguese man of war are close to shore, where news stories warn of purple flags, vicious stings, and ruined beach days, however we still know almost nothing about their behavior, ecology, and lifecycle out in the open ocean.

## Methods

### Collecting and fixing

Juvenile specimens, defined as colonies with float length 8-10cm and immature developing gonodendra, were collected from locations along the exposed Gulf coast of Galveston Island, TX in March 2016 and February 2017, from East Beach (Lat. Lon. 29.328090, −94.737542) to east of Galveston Island State park (Lat. Lon. 29.195358, −94.948335). Information on when to collect large numbers of *P. physalis* was obtained from sightings submitted to the citizen science website Jelly Watch (www.jellywatch.org). Juvenile specimens were collected alive fresh from the surf, and transferred directly to the lab for examination and fixation in 4% formalin in seawater after relaxation in 7.5% MgCl_2_ hexahydrate in distilled water, mixed 1/3 with seawater. Physical vouchers are deposited at the Peabody Museum of Natural History (Yale University), New Haven, CT (exemplar specimens YPM 103765, YPM 103766). Developing specimens were obtained from the collections of Philip R. Pugh, and are now deposited at the Peabody Museum of Natural History (Yale University). These specimens were collected in various locations in the Atlantic Ocean during research expeditions in 1972, 1973 and 1983. Two additional vials of developing specimens were kindly provided by Dr. Pugh, however no collection information is available. Details of the collected specimens are provided in table 1.

**Table 1:** Collection information for larval developing Portuguese man of war specimens used in this study

| Date of collection | Latitude, Longitude | Depth (m) | Expedition information | Catalog number |
| --- | --- | --- | --- | --- |
| 18 February 1972 | 22.80555556, -22.55277778 | 0 | RRS Discovery 7800 | YPM 105036 |
| 22 February 1972 | 17.93888889, -24.88611111 | 0 | RRS Discovery 7803 #13 | YPM 105035 |
| 1 March 1973 | 32.00277778, -34.38055556 | 0 | RRS Discovery 8270 | YPM 105037 |
| 1 August 1983 | 13.31861111, -56.01861111 | NA | BWP 1093-15 PPP106 | YPM 105038 |

### Image capture and processing

Optical Projection Tomography (OPT) was used as a tool to collect serial images for three-dimensional reconstruction of fixed *P. physalis* tissue. Before imaging, formalin fixed specimens were washed 2x quickly in cold phosphate buffered saline (PBS), 3x PBS (5 min) and placed in 1:1000 4’,6-diamidino-2-phenylindole (DAPI) and PBS overnight to fluorescently stain nuclei. Samples were then washed in PBS twice (5 min) and embedded in 2.5% Ultra Low Melting Point Agarose (Invitrogen, 16520-050) within a syringe cylinder. Agarose cylinders, containing the embedded tissue, were dehydrated into methanol (25% MeOH/H_2_O, 50% MeOH/H_2_O, 75% MeOH/H_2_O for 4hs each step, and 100% MeOH overnight). Specimens were optically cleared using benzyl alcohol-benzyl benzoate (BABB) (MeOH/BABB (1:1) 1h, BABB overnight). Specimens were then imaged on a custom built optical projection tomography system in the Optical Imaging & Vital Microscopy Core, Baylor College of Medicine, Houston TX, with a camera pixel size of 6.7um, an image pixel size of 8.75um, and a round scanning trajectory. OPT images were reconstructed using NRecon software (Bruker microCT, v. 1.3). The files were subsequently resampled for segmentation and volume rendering by removing every other slice and also by scaling the images by half. The 3D reconstructions were created and segmented using Amira Software (ThermoScientific v. 5.3.3). Raw fluorescent images are included in supplementary information (Figs. S1, S2, S3, S4).

## Acknowledgements

CM would like to thank citizen scientists and jellywatchers Lea, Millie, Dignan, and Noah for alerting her, via Jellywatch (http://jellywatch.org/), to the presence of large numbers of Portuguese man o’ war along the coast in 2016, and for their help collecting specimens. CM would also like to thank Maria Pia Miglietta and her lab members at Texas A&M Galveston for hosting her during field visits. We would also like to thank Phil Pugh for his thorough and helpful feedback on this manuscript, and Zack Lewis for help with Amira. This work was supported by Marine Biological Laboratory Embryology post-course research funds, an NSF Doctoral Dissertation Improvement Grant (DDIG) in the Directorate for Biological Sciences (NSF DEB-1701272), and NSF DEB-1256695.

## References

1. Zaitsev, Y. Neuston of seas and oceans. in The sea surface and global change (eds. Liss, P. S. & Duce, R. A.) 371–382 (Cambridge University Press, 1997).

2. Clark, F. E. & Lane, C. E. Composition of float gases of *Physalia physalis*. Proceedings of the Society for Experimental Biology and Medicine 107, 673–674 (1961).

3. Iosilevskii, G. & Weihs, D. Hydrodynamics of sailing of the Portuguese man-of-war *Physalia physalis*. Journal of the Royal Society Interface 6, 613–626 (2009).

4. Prieto, L., Macías, D., Peliz, A. & Ruiz, J. Portuguese Man-of-War (*Physalia physalis*) in the Mediterranean: A permanent invasion or a casual appearance? Scientific reports 5, 11545 (2015).

5. Mapstone, G. M. Global diversity and review of Siphonophorae (Cnidaria: Hydrozoa). PLoS One 9, e87737 (2014).

6. Munro, C. et al. Improved phylogenetic resolution within Siphonophora (Cnidaria) with implications for trait evolution. Molecular Phylogenetics and Evolution 127, 823–833 (2018).

7. Pugh, P. R. The diel migrations and distributions within a mesopelagic community in the north east Atlantic. 7. Siphonophores. Prog. Oceanogr. 13, 461–489 (1984).

8. Dunn, C. W., Pugh, P. R. & Haddock, S. H. D. Molecular Phylogenetics of the Siphonophora (Cnidaria), with Implications for the Evolution of Functional Specialization. Syst. Biol. 54, 916–935 (2005).

9. Mackie, G. O., Pugh, P. R. & Purcell, J. E. Siphonophore biology. Advances in Marine Biology 24, 97–262 (1987).

10. Araya, J. F., Aliaga, J. A. & Araya, M. E. On the distribution of *Physalia physalis*(Hydrozoa: Physaliidae) in Chile. Marine Biodiversity 46, 731–735 (2016).

11. Copeland, D. E. Fine Structures Of The Carbon Monoxide Secreting Tissue In The Float Of Portuguese Man-of-war *Physalia physalis* L. The Biological Bulletin 135, 486–500 (1968).

12. Herring, P. J. Biliprotein coloration of *Physalia physalis*. Comparative Biochemistry and Physiology Part B: Comparative Biochemistry 39, 739–746 (1971).

13. Lane, C. E. The toxin of *Physalia* nematocysts. Annals of the New York Academy of Sciences 90, 742–750 (1960).

14. Larimer, J. L. & Ashby, E. A. Float gases, gas secretion and tissue respiration in the Portuguese man-of-war, *Physalia*. Journal of Cellular and Comparative Physiology 60, 41–47 (1962).

15. Totton, A. K. & Mackie, G. O. Dimorphism in the Portuguese man-of-war. Nature 177, 290 (1956).

16. Wilson, D. P. The Portuguese Man-of-War, *Physalia physalis*L., in British and adjacent seas. Journal of the Marine Biological Association of the United Kingdom 27, 139–172 (1947).

17. Wittenberg, J. B. The source of carbon monoxide in the float of the Portuguese man-of-war, *Physalia physalis* L. Journal of Experimental Biology 37, 698–705 (1960).

18. Wittenberg, J. B., Noronha, J. M. & Silverman, M. Folic acid derivatives in the gas gland of *Physalia physalis* L. Biochemical Journal 85, 9 (1962).

19. Woodcock, A. H. Dimorphism in the Portuguese man-of-war. Nature 178, 253 (1956).

20. Yanagihara, A. A., Kuroiwa, J. M. Y., Oliver, L. M. & Kunkel, D. D. The ultrastructure of nematocysts from the fishing tentacle of the Hawaiian bluebottle, *Physalia utriculus* (Cnidaria, Hydrozoa, Siphonophora). Hydrobiologia 489, 139–150 (2002).

21. Bardi, J. & Marques, A. C. Taxonomic redescription of the Portuguese man-of-war, *Physalia physalis* (Cnidaria, Hydrozoa, Siphonophorae, Cystonectae) from Brazil. Iheringia. Ser. Zool. 97, 425–433 (2007).

22. Mackie, G. O. Studies on *Physalia physalis* (L.). Part 2. Behavior and histology. Discovery Reports 30, 371–407 (1960).

23. Okada, Y. K. Développement post-embryonnaire de la Physalie Pacifique. Memoirs of the College of Science, Kyoto Imperial University, Series B 8, 1–27 (1932).

24. Okada, Y. K. Les jeunes Physalies. Memoirs of the College of Science, Kyoto Imperial University, Series B 10, 407–410 (1935).

25. Steche, O. Die knospungsgesetz und der Bau der Anhangsgruppen von Physalia. in Festschrift zum sechzigsten geburtstag richard hertwigs 356–372 (Verlag Von Gustave Fischer, 1910).

26. Totton, A. K. Studies on *Physalia physalis* (L.). Part 1. Natural history and morphology. Discovery Reports 30, 301–368 (1960).

27. Totton, A. K. A synopsis of the Siphonophora. (British Museum (Natural History), 1965).

28. Dunn, C. W. & Wagner, G. P. The evolution of colony-level development in the Siphonophora (Cnidaria: Hydrozoa). Dev. Genes Evol. 216, 743–754 (2006).

29. Church, S. H., Siebert, S., Bhattacharyya, P. & Dunn, C. W. The histology of *nanomia bijuga* (hydrozoa: Siphonophora). Journal of Experimental Zoology Part B: Molecular and Developmental Evolution 324, 435–449 (2015).

30. Jenkins, R. L. Observations on the commensal relationship of *Nomeus gronovii* with *Physalia physalis*. Copeia 1983, 250–252 (1983).

31. Cartwright, P., Bowsher, J. & Buss, L. W. Expression of a Hox gene, Cnox-2, and the division of labor in a colonial hydroid. Proceedings of the National Academy of Sciences 96, 2183–2186 (1999).

32. Cartwright, P. & Nawrocki, A. M. Character evolution in Hydrozoa (phylum Cnidaria). Integrative and Comparative Biology 50, 456–472 (2010).

33. Schuchert, P. Hydroids (Cnidaria, Hydrozoa) of the Danish expedition to the Kei Islands. Steenstrupia 27, 137–256 (2003).

34. Haeckel, E. Report on the Siphonophorae collected by HMS Challenger during the years 1873-1876. Report of the Scientific Results of the voyage of H.M.S. Challenger. Zoology 28, 1–380 (1888).

35. Huxley, T. H. The Oceanic Hydrozoa: a description of the Calycophoridae and Physophoridae observed during the voyage of HMS Rattlesnake 1846-1850. (The Ray Society, 1859).

36. Haddock, S. H. D., Dunn, C. W. & Pugh, P. R. A re-examination of siphonophore terminology and morphology, applied to the description of two new prayine species with remarkable bio-optical properties. J. Mar. Biol. Assoc. U.K. 85, 695–707 (2005).

37. Carré, D. Etude histologique du developpement de *Nanomia bijuga* (Chiaje, 1841), Siphonophore Physonecte, Agalmidae. Cah. Biol. Mar. 10, 325–341 (1969).

38. Agassiz, A. & Mayer, A. G. Reports on the scientific results of the expedition to the Tropical Pacific, 1899-1900. III. The Medusae. Memoirs of the Museum of Comparative Zoology at Harvard University 26, 139–176 (1902).

39. Levin, M. Left–right asymmetry in embryonic development: a comprehensive review. Mechanisms of development 122, 3–25 (2005).

40. Dunn, C. W. Complex colony-level organization of the deep-sea siphonophore *Bargmannia elongata* (Cnidaria, Hydrozoa) is directionally asymmetric and arises by the subdivision of pro-buds. Developmental dynamics 234, 835–845 (2005).

41. Siebert, S. et al. Stem cells in *Nanomia bijuga* (Siphonophora), a colonial animal with localized growth zones. EvoDevo 6, 22 (2015).

42. Siebert, S., Pugh, P. R., Haddock, S. H. D. & Dunn, C. W. Re-evaluation of characters in Apolemiidae (Siphonophora), with description of two new species from Monterey Bay, California. Zootaxa 3702, 201–232 (2013).

43. Pickwell, G. V., Barham, E. G. & Wilton, J. W. Carbon monoxide production by a bathypelagic siphonophore. Science 144, 860–862 (1964).

44. Chun, C. Zur Morphologie der Siphonophoren. 2. Ueber die postembryonale Entwickelung von *Physalia*. Zoologischer Anzeiger 10, 557–577 (1887).

45. Lamarck, J. B. Système des animaux sans vertèbres. (Deterville, 1801).

46. Pontin, D. R. & Cruickshank, R. H. Molecular phylogenetics of the genus *Physalia* (Cnidaria: Siphonophora) in New Zealand coastal waters reveals cryptic diversity. Hydrobiologia 686, 91–105 (2012).

47. Woodcock, A. H. A theory of surface water motion deduced from the wind-induced motion of the *Physalia*. J. mar. Res 5, 196–206 (1944).

48. Clark, A. M. The Portuguese Man-of-War. Marine Observer 40, 63–65 (1970).

49. Purcell, J. E. Feeding ecology of *Rhizophysa eysenhardti*, a siphonophore predator of fish larvae. Limnology and Oceanography 26, 424–432 (1981).

50. Purcell, J. E. Predation on fish larvae by *Physalia physalis*, the Portuguese man of war. Marine ecology progress series. Oldendorf 19, 189–191 (1984).

51. Lane, C. E. & Dodge, E. The toxicity of *Physalia* nematocysts. The Biological Bulletin 115, 219–226 (1958).

52. Lenhoff, H. M. & Schneiderman, H. A. The chemical control of feeding in the Portuguese man-of-war, *Physalia physalis* L. and its bearing on the evolution of the Cnidaria. The Biological Bulletin 116, 452–460 (1959).

53. Mackie, G. O. & Boag, D. A. Fishing, feeding and digestion in siphonophores. Pubbl statz zool Napoli 33, 178–96 (1963).

54. Babcock, H. L. The Sea-Turtles of the Bermuda Islands, with a Survey of the present state of the Turtle Fishing Industry. in Proceedings of the zoological society of london 107, 595–602 (Wiley Online Library, 1938).

55. Bingham, F. O. & Albertson, H. D. Observations on beach strandings of the *Physalia* (Portuguese-man-of-war) community. Veliger 17, 220–224 (1974).

56. Bieri, R. Feeding preferences and rates of the snail, *Ianthina prolongata*, the barnacle, *Lepas anserifera*, the nudibranchs, *Glaucus atlanticus* and *Fiona pinnata*, and the food web in the marine neuston. Publ. Seto Mar. Biol. Lab. 14, 161–170 (1966).

57. Thompson, T. E. & Bennett, I. *Physalia* nematocysts: utilized by mollusks for defense. Science 166, 1532–1533 (1969).

58. Valdés, Á. & Campillo, O. A. Systematics of pelagic aeolid nudibranchs of the family Glaucidae (Mollusca, Gastropoda). Bulletin of Marine Science 75, 381–389 (2004).

59. Kato, K. Is *Nomeus* a harmless Inquilinus of *Physalia*? Proceedings of the Imperial Academy 9, 537–538 (1933).

60. Lane, C. E. The Portuguese man-of-war. Scientific American 202, 158–171 (1960).

