## Supplementary Figures for "Morphology and development of the Portuguese man of war, *Physalia physalis*"

### Supplementary Information

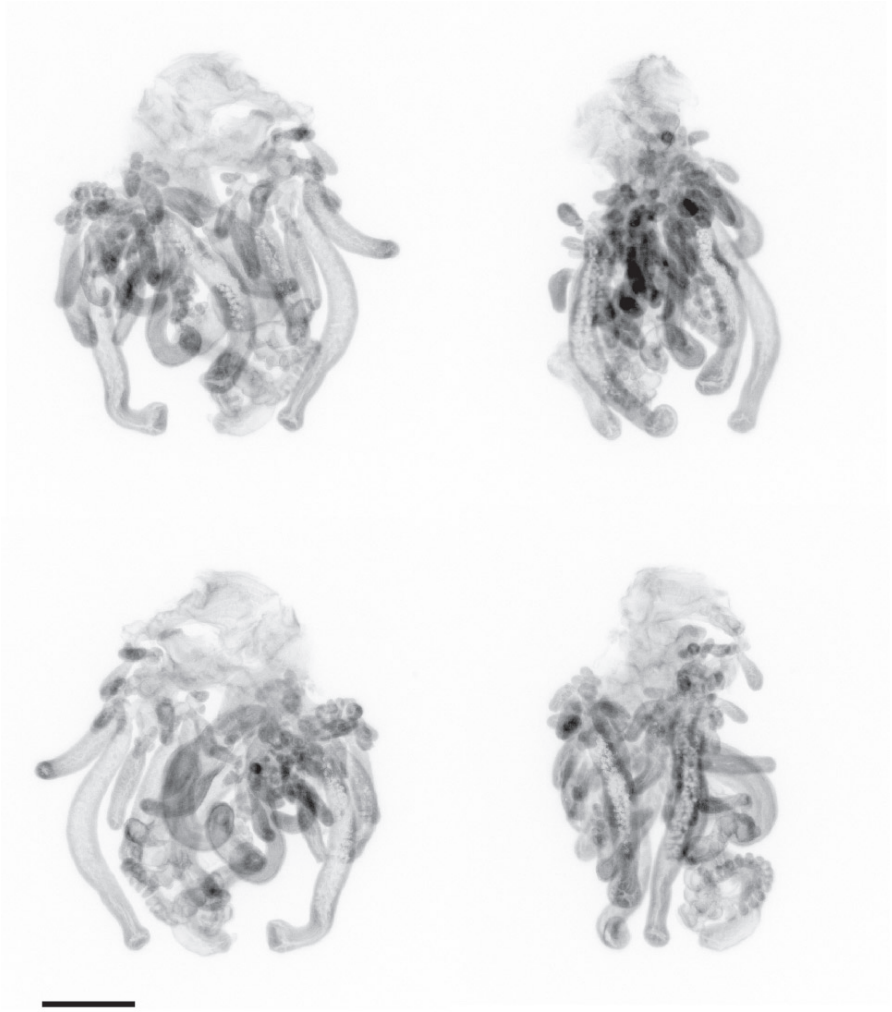

Figure S1: Raw fluorescent images of formalin fixed juvenile *Physalia physalis* zooids, images obtained by optical projection tomography. Images are different views of the same specimen, the same specimen as in fig. 7. Scale bar is 2mm.

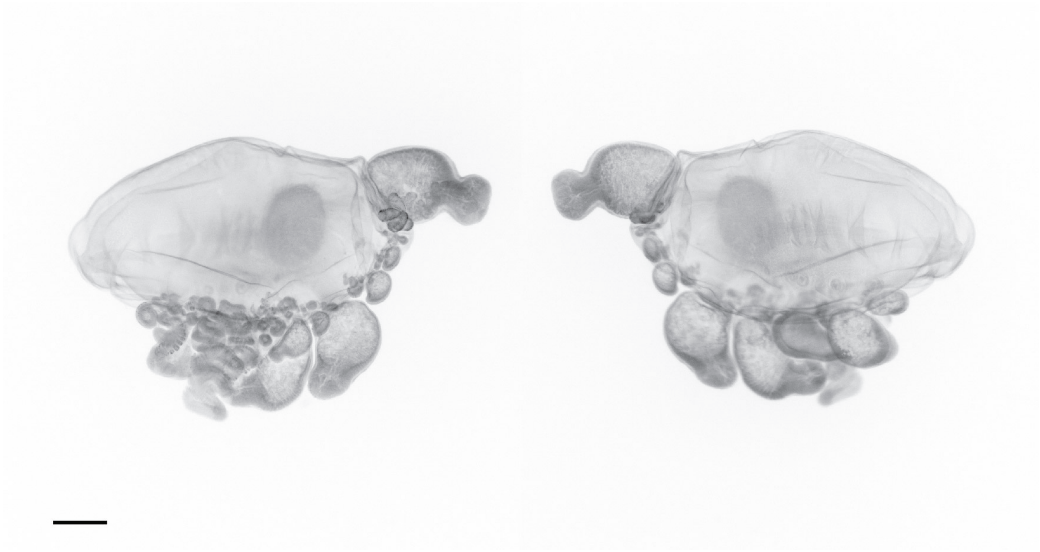

Figure S2: Raw fluorescent images of formalin fixed larval *Physalia physalis*, images obtained by optical projection tomography. Images are different views of the same specimen, the same specimen as in fig. 4. Scale bar is 1mm.

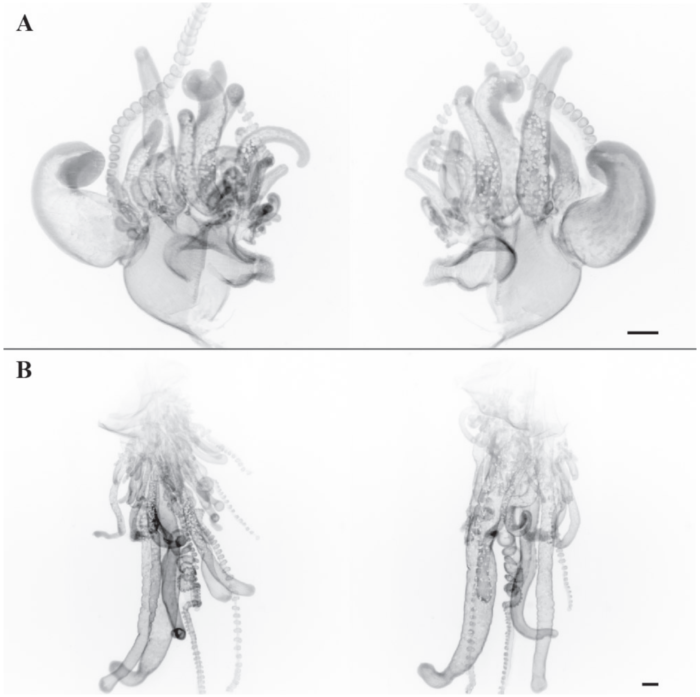


Figure S3: Raw fluorescent images of formalin fixed juvenile *Physalia physalis* zooids from the same specimen, images obtained by optical projection tomography. Scale bar is 1mm. A. The posterior zone including protozooid as the posterior most zoid. B. Section including mature gastrozooids, tentacular palpons, and developing tripartite groups.

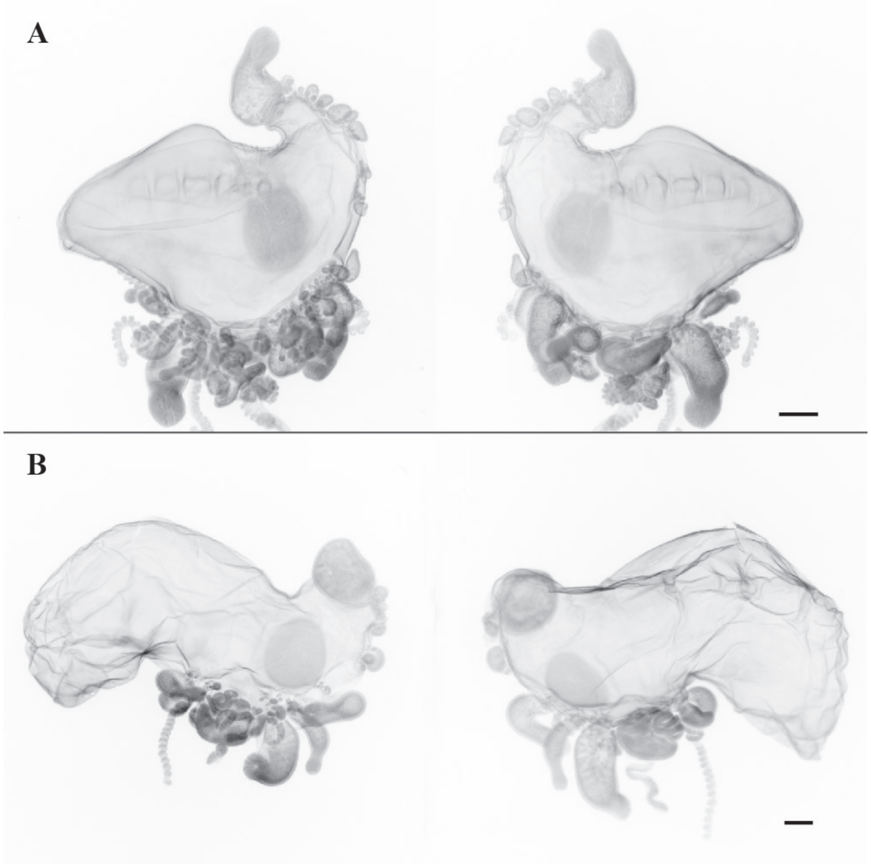


Figure S4: Raw fluorescent images of formalin fixed larval *Physalia physalis*, images obtained by optical projection tomography. Scale bar is 1mm. A & B are different specimens.

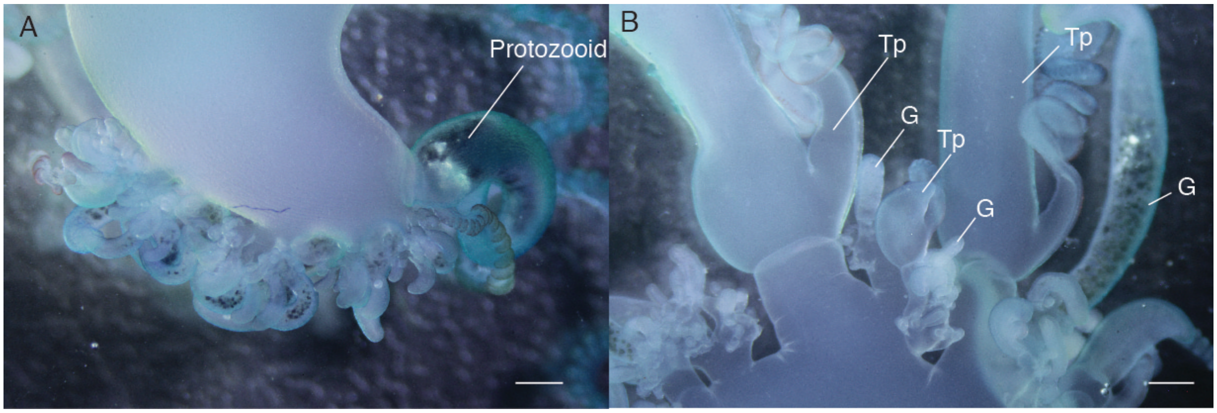


Figure S5: Regions of growth in juvenile *Physalia physalis* (float length 8-10 cm). A. Photograph of the posterior zone, with protozooid as the posterior most zooid. B. Photograph of mature tentacular palpons and gastrozooids, and developing tripartite groups forming proximally. Tp: Tentacular palpon; G: Gastrozooid. Scale bar is 1 mm.
